# Determination of Gaussian curvature modulus and spontaneous curvature via membrane buckling

**DOI:** 10.1101/2024.12.27.630512

**Authors:** Mei-Ting Wang, Gao-Xiao Jiang, Rui Ma, Chen-Xu Wu

## Abstract

The elastic properties of membranes are typically characterized by a few phenomenological parameters, including bending and Gaussian curvature moduli measuring the membrane rigidity against its deformation and topological change, as well as spontaneous curvature arising from the asymmetry between the two leaflets in the lipid bilayers. Though tether-based and fluctuation-based experiments are commonly utilized to measure the bending modulus, measuring the Gaussian curvature modulus and the spontaneous curvature of the membrane is considered to be much more difficult. In this paper, we study the buckling process of a circular membrane with nonzero spontaneous curvature under compressive stresses. It is found that when the stress exceeds a critical value, the circular membrane will transform from a spherical cap to a buckled shape, with its buckling degree enhanced with the increase of stress until its base is constricted to almost zero. As the stress-strain relationship of the buckled membrane strongly depends on the Gaussian curvature modulus and the spontaneous curvature,we therefore propose a method to determine the Gaussian curvature modulus and the spontaneous curvature simultaneously by measuring its stress-strain relationship during a buckling process.

## I. INTRODUCTION

The mechanical properties of membranes play a crucial role in their function, structure, and dynamics in biological systems [1]. These properties are primarily governed by the composition and structure of the lipid bilayer, and its interactions with proteins. In 1973, by making analogies between the lipid bilayer and the liquid crystal in their orientational order, Helfrich proposed an elastic theory for the cell membrane [2]. As the thickness of the membrane is much smaller than its lateral dimension, the theory assumes that the membrane is a two-dimensional surface embedded in three-dimensional space, and deforming under forces while maintaining its in-plane fluidity [2, 3]. By writing the bending energy of a membrane in terms of the mean curvature and the Gaussian curvature of the membrane surface, Helfrich theory has been successfully applied to describing and predicting various deformation phenomena of cell membranes, including the biconcave shape of the red blood cells [4, 5], vesicle shape transformation [6–11], membrane fusion, fission [12–15], as well as membrane-protein interaction [16–22].

In Helfrich theory, the mechanical properties of membrane are characterized by three phenomenological parameters: the bending modulus *κ*, the Gaussian curvature modulus 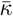, and the spontaneous curvature *c*_0_. The equilibrium membrane shape is obtained by minimizing the elastic energy subject to geometric constraints. The bending modulus *κ* quantifies the energetic cost of deforming the membrane’s mean curvature from its spontaneous curvature *c*_0_. Any deformation that induce membrane curvature could be related with *κ*, therefore, experimental determination of the bending modulus *κ* can be achieved by probing the mechanical response of the membrane to forces [23–27] or thermal fluctuations [28–33]. For instance, measuring the force required to pull a membrane tether from a giant vesicle [25–27] or the fluctuation spectrum of a freely standing membrane [31–33] is widely used to obtain the value of the bending modulus.

Spontaneous curvature *c*_0_ reflects the membrane’s tendency to adopt a preferred curvature even in the absence of external forces. Any types of asymmetry between the two leaflets, such as the lipid composition, lipid number density, and protein occupation, could induce spontaneous curvature [34–37]. Directly measuring the spontaneous curvature of a membrane in experiments is challenging, as its effect is always intertwined with the bending modulus *κ* and the membrane tension. However, a comparison between the experimentally observed membrane shape and the theory-predicted membrane shape in response to an asymmetric factor provides an indirect approach to extract the spontaneous curvature.

Unlike the bending modulus *κ* governing smooth deformations, the Gaussian bending modulus 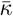 contributes to the total energy only in processes involving changes in topology (e.g., transitions from a sphere to a torus, formation of pores, vesicle budding, or fission events) due to the famous Gauss-Bonnet theorem [38, 39]. This makes direct measurement of 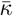 less straightforward. As the formation and closure of membrane pores involve Gaussian curvature contributions [40], measuring the force to close the pore [41] or reading the fluctuations at the edge of an open membrane [42] has been used to determine the Gaussian curvature modulus 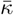 in molecular dynamics simulations.

The buckling phenomenon is a mechanical instability that occurs when a structure subject to compressive stress undergoes a sudden deformation or collapse [43]. In various biological processes, such as the spiky shape of red blood cells induced by osmotic stress [44], or the formation of filopodia during cell migration [45], such membrane buckling is associated with a creation of surface folds or wrinkles. Here it should be noted that buckling of a fluid membrane exhibits different behaviors compared with that of a solid sheet, such as anisotropic stress and negative compressibilities [46, 47]. This has been used as a robust method to extract the bending modulus *κ* of a rectangular membrane when subject to compressive stresses [47].

In our previous work, we proposed a method to measure the Gaussian curvature modulus 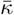 of a circular membrane via a buckling protocol [48]. The membrane is subject to radially oriented compressive stresses and the critical stress to buckled the membrane is found to be dependent on 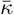 if no external torque is applied at the edge of the membrane. However, the effect of the spontaneous curvature is not considered in this study. In this work, we extend our previous work to conditions where the buckled membrane possesses a nonzero spontaneous curvature. It is shown that the stress-strain relationship of a buckled membrane depends on both the Gaussian curvature modulus 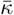 and the spontaneous curvature *c*_0_. Based on this observation, we propose a method capable of measuring 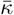 and *c*_0_ simultaneously.

## II. THEORETICAL MODEL

The membrane in this paper is modelled as a twodimensional axisymmetric surface parameterized as

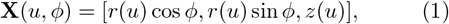

where the parameter *ϕ* ∈ [0, 2*π*] and *u* ∈ [0, 1], with *u* = 0 and *u* = 1 corresponding to the membrane tip and the membrane base, respectively (Fig. 1). The shape functions *r*(*u*) and *z*(*u*) satisfy

**FIG. 1.**
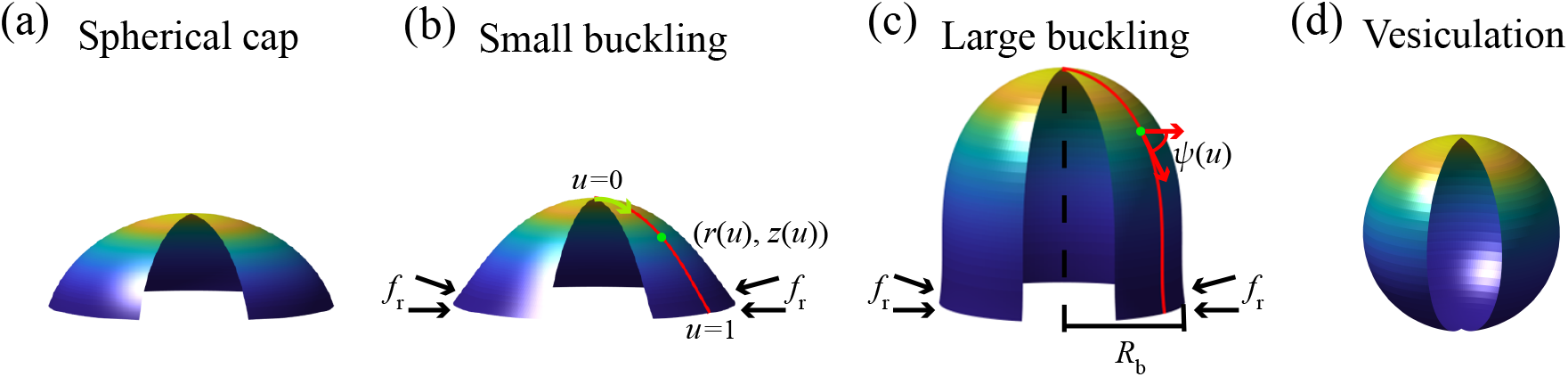
Illustration of deformed circular membranes with (a) a spherical cap, (b) a small buckling, and (c) a large buckling or (d) a vesiculation under free-hinge BC, considering spontaneous curvature.

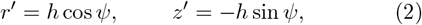

where *ψ*(*u*) represents the angle spanned between the tangential direction of the meridian curve **e**_*u*_ = *∂***X***/∂u* and the radial direction **e**_*r*_ = [cos *ϕ*, sin *ϕ*, 0],*h*(*u*) is a scaling factor that fulfills the equation 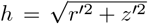 Hereafter, we use the prime to denote the derivative with respect to *u*. Note that when the parameter *u* varies from 0 to 1, the two coordinate expressions [*r*(*u*), *z*(*u*)] and [*r*(*u*^2^), *z*(*u*^2^)] describe the same membrane shape but differ in the expression of *h*. Due to this redundancy, we impose *h*^′^ = 0, indicating that *h* is a constant, which is equal to the total arclength of the membrane profile.

For a membrane with a uniform spontaneous curvature *c*_0_, its bending energy can be written as

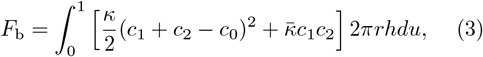

where *κ* and 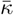 denote the bending rigidity of the membrane associated with the mean curvature and the Gaussian curvature, respectively, *c*_1_ = *ψ*^′^*/h* and *c*_2_ = (sin *ψ*)*/r* are the two principal curvatures of the membrane surface.

To account for the compressive stress exerted at the membrane base, we introduce a boundary energy term

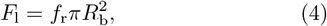

where *f*_r_ denotes the magnitude of the compressive stress pointing inwards along the radial direction, and *R*_b_ denotes the radius of the membrane base. The total free energy *F* of the membrane includes the bending energy and the boundary energy,

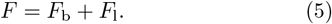

We assume that the membrane is incompressible, which means that the total surface area *A*_0_ of the membrane is conserved when deformed. Such a constraint is imposed by introducing a Lagrangian multiplier *σ* conjugated to the surface area *A*,

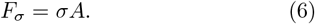

The effective energy of the membrane 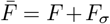 can be expressed as a functional of the shape variables for the meridian profile:

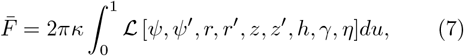

where *γ* and *η* are Lagrangian multipliers that enforce the geometric constraints for Eq. (2). A variation of the functional 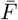 with respect to the shape variables leads to shape equations (A3), (A4) and (A5), as well as boundary conditions (BCs) Eqs. (B1)-(B6) and (B9), forming a boundary value problem (see Appendix A and B for the mathematical details). In general, there are two types of BCs, depending on whether the angle at the membrane base is free to rotate (free-hinge BC) or fixed to a constant value (fixed-hinge BC). As we have shown previously that only with the free-hinge BC will the buckling process depend on the Gaussian bending rigidity 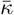 [48], we restrain our investigation to the free-hinge BC, which is given by

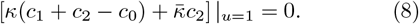

When the membrane is in its native state without any external forces applied, it tends to form a spherical cap with a radius of

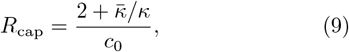

which is obtained from the BC of Eq. (8), given that a spherical cap has *c*_1_ = *c*_2_ = 1*/R*_cap_ (Fig. 1a). The spanning angle *θ* of the spherical cap is determined by its surface area *A*_0_ via the equation

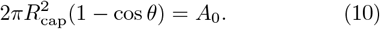

The bending energy of such a spherical cap reads

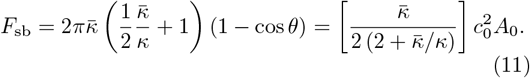

Not e that the spherical cap shape expressed in Eqs. (9) and (10) is always a solution to the shape equations (A3)-(A5), even when a compressive stress *f*_r_ is applied at the base perimeter of the membrane. However, the total free energy of the spherical cap solution

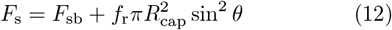

increases linearly with the stress *f*_r_ due to the boundary energy term Eq. (4).

## III. RESULTS AND DISCUSSIONS

### A. Buckling of membranes with different Gaussian curvature moduli

In this section, we study the effects of Gaussian curvature modulus on the buckling of a circular membrane in response to a compressive stress *f*_r_. The spontaneous curvature of the membrane is fixed at *c*_0_*R*_0_ = 0.1. In addition to the spherical cap solution ( Fig. 2, cyan line), a new branch of solutions, which represent a buckled membrane shape, emerges when the stress *f*_r_ is beyond a critical value 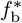 (black curve in Fig. 2). The total free energy of the buckled membrane shape *F* = *F*_b_ +*F*_l_ is lower than that of the spherical cap *F*_s_, indicating the occurrence of a buckling transition. Upon further increase of the stress *f*_r_ to 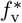, the degree of membrane buckling is enhanced until the base radius is narrowed down to almost zero (Fig. 2(d)), which is termed vesiculation for the rest of the paper. When comparing the free energy curves for different Gaussian curvature moduli 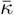, we find that both the critical stress of membrane buckling 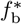 and the stress of vesiculation 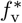 are reduced for more negative 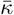 (compare Fig. 2a, b, and c). In particular, when 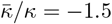, the buckled shape solutions appear even in the absence of compressive stresses and has a lower free energy than the spherical cap solution, indicating a spontaneous buckling of the membrane (Fig. 2c).

**FIG. 2.**
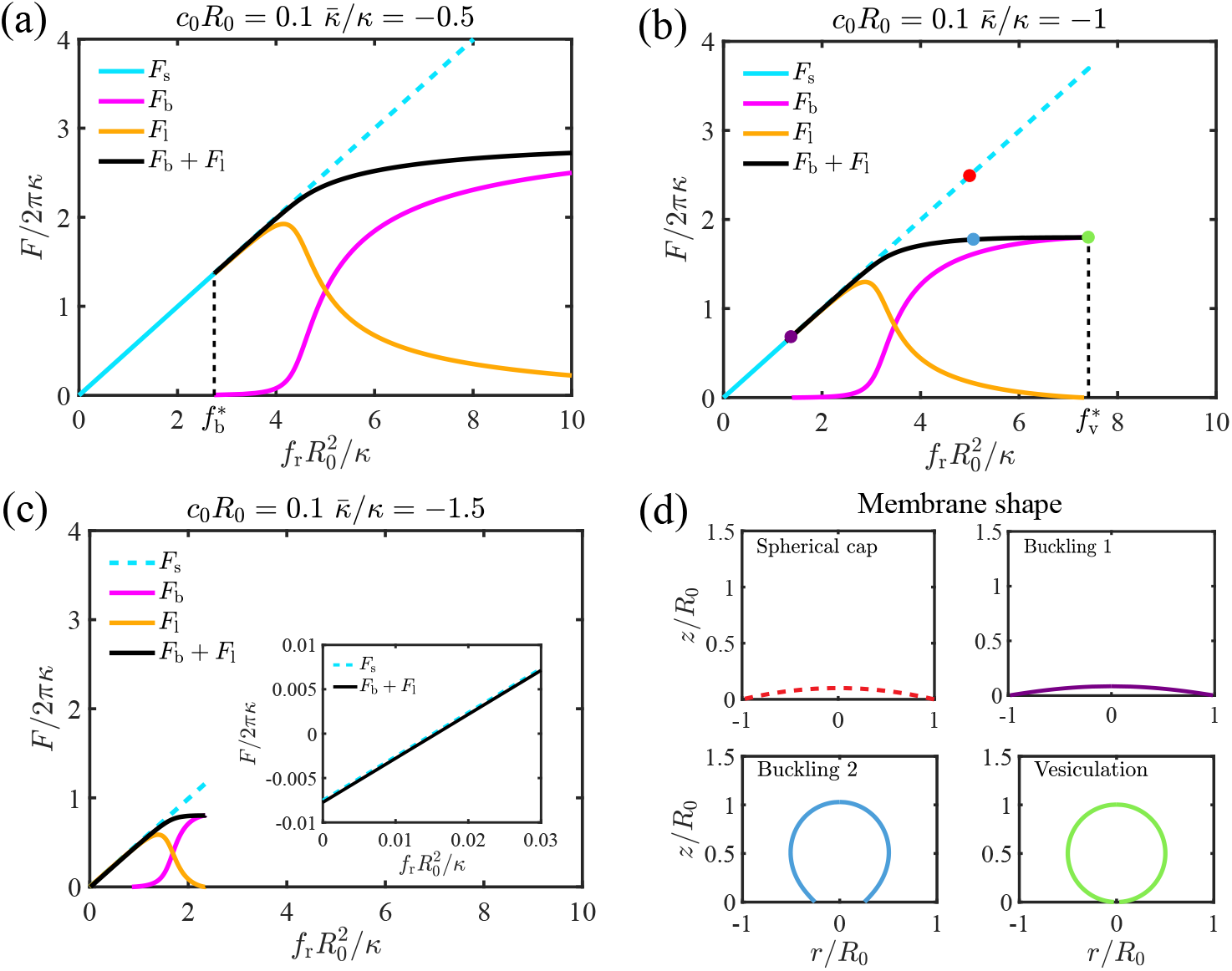
Effect of the stress 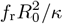 on the free energy *F/*2*πκ* of the membrane with a scaled spontaneous curvature *c*_0_*R*_0_ = 0.1 based on different Gaussian curvatures (a) 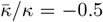 (b) 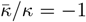, and (c) 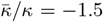. (d) demonstrates the membrane shapes corresponding to the colored dots in (b).

To systematically investigate how the Gaussian curvature moduli 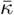 influence the buckling behavior of a circular membrane, we show the phase diagram of the membrane shape as a function of 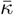 and the stress *f*_r_ in Fig. 3 for different spontaneous curvature *c*_0_. When *c*_0_*R*_0_ = 0.1 or 0.2, the phase diagram is divided into three parts, separated by the curve of critical buckling stress 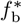 (Fig. 3a and b, the boundary between the pink region and the grey region), and the curve of critical vesiculation stress 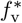 (Fig. 3a and b, the boundary between the pink region and the blue region). The area of the grey region, which represents the spherical cap shape, is reduced with increasing spontaneous curvature *c*_0_, and vanishes when *c*_0_ is large (Fig. 3c). Overall, the results suggest that a negative Gaussian curvature modulus 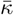 and a positive spontaneous curvature *c*_0_ tend to facilitate the buckling by reducing the critical buckling stress 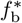. However, the critical vesiculation stress 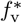 is not strongly influenced by the spontaneous curvature.

**FIG. 3.**
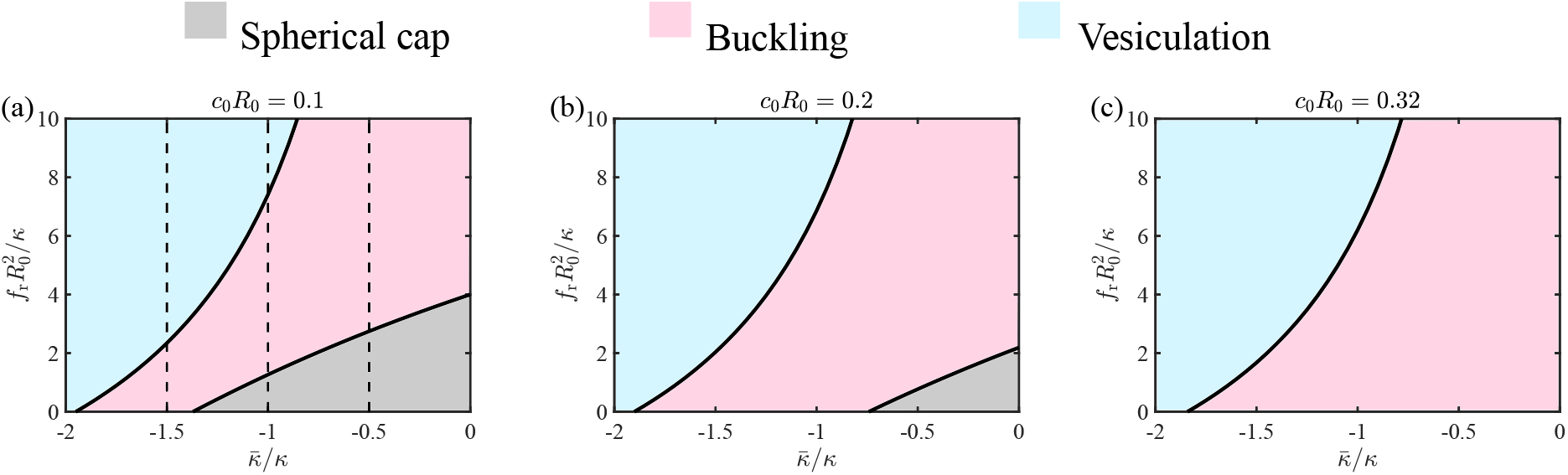
Phase diagram of the membrane shape as a function of the Gaussian curvature modulus 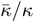 and the compressive stress 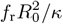 under different spontaneous curvatures (a) *c*_0_*R*_0_ = 0.1, (b) *c*_0_*R*_0_ = 0.2 and (c) *c*_0_*R*_0_ = 0.32. The dashed lines in (a) represent the cases of Fig. 2(a), (b) and (c), respectively.

The effect of the Gaussian curvature modulus on the buckling behavior of membrane can be further reflected on the stress-strain relationship shown in Fig. 4. We quantify the buckling degree of the membrane by introducing the strain *μ*_r_ = (*A*_0_ −*A*_b_)*/A*_0_, where 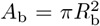 represents the projected area of the buckled membrane. The stress *f*_r_(*μ*_r_ = 0) defines the critical buckling stress 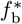 and the stress *f*_r_(*μ*_r_ = 1) defines the critical vesiculation stress 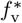. At the beginning of the buckling when *μ*_r_ is small, the stress *f*_r_ increases rapidly with *μ*_r_. At intermediate *μ*_r_, the stress *f*_r_ mildly with *μ*_r_. At the end of the buckling when *μ*_r_ is close to 1, the stress *f*_r_ rises rapidly again with *μ*_r_. The vesiculation stress 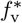 can be significantly reduced with more negative 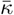.

**FIG. 4.**
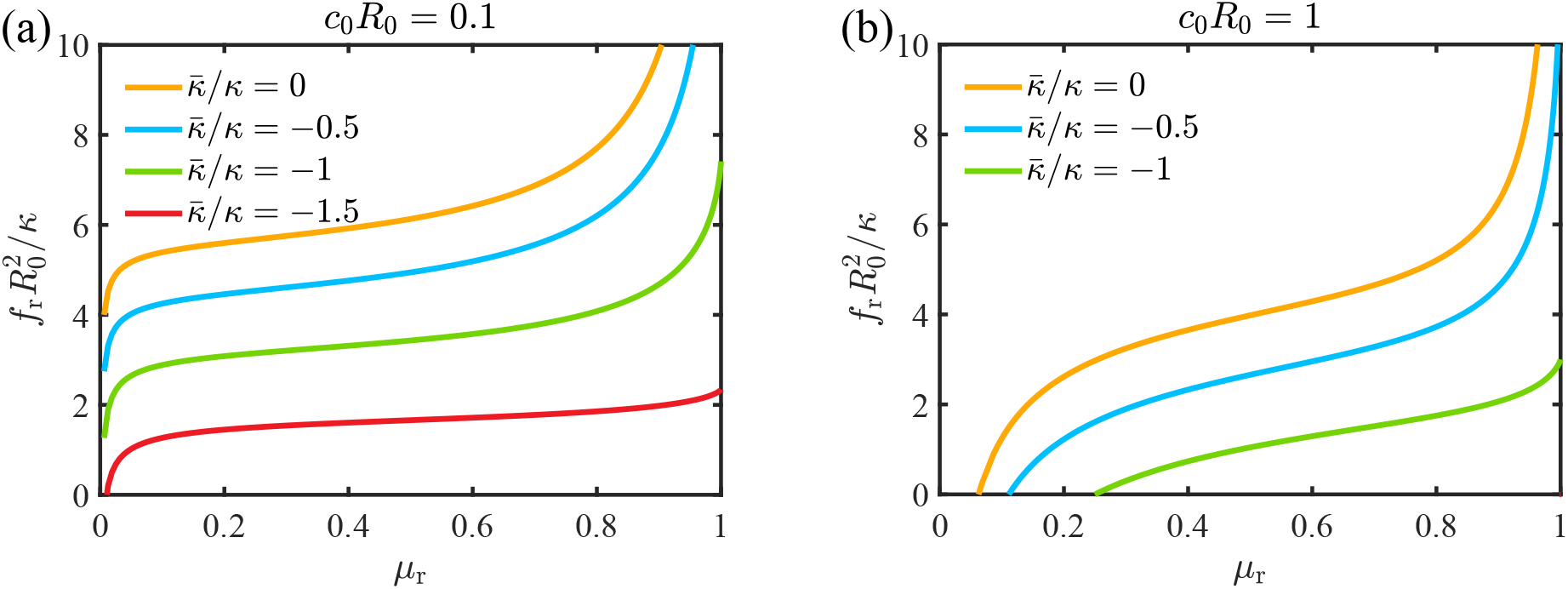
The stress-strain relationship of the buckled membrane shape under different Gaussian curvature moduli, with the spontaneous curvatures (a) *c*_0_*R*_0_ = 0.1 and (b) *c*_0_*R*_0_ = 1.

### B. Buckling of membranes with different spontaneous curvatures

In this section, we investigate the buckling behavior of a circular membrane under different spontaneous curvatures *c*_0_*R*_0_. The phase diagrams of the membrane shape as a function of the compressive stress *f*_r_ and the spontaneous curvature *c*_0_ are shown in Fig. 5 for 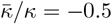 and 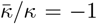, respectively. It is found that increasing the spontaneous curvature *c*_0_ is able to reduce the critical buckling stress 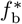, which is the boundary that separates the grey region from the pink region. When *c*_0_ exceeds a critical value, the critical buckling stress 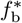 turns negative (Fig. 5), which implies spontaneous buckling in the absence of compressive stresses. The vesiculation buckling stress 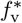, which is the boundary that separates the blue region from the pink region, is also reduced with increasing *c*_0_ (Fig. 5). The reduction is more dramatic for more negative Gaussian curvature modulus. spontaneous curvatures as shown in Fig. 5.

**FIG. 5.**
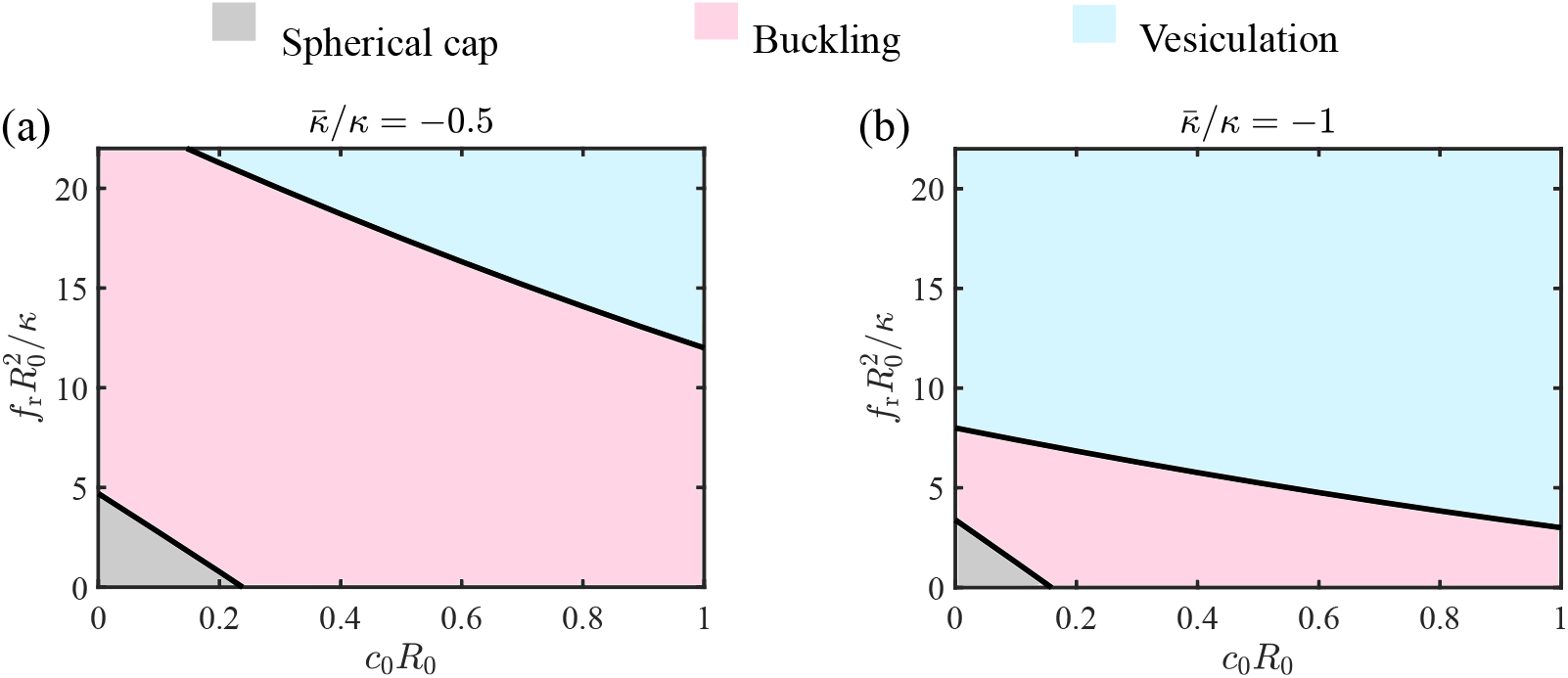
Phase diagram of the membrane constructed by the scaled spontaneous curvature *c*_0_*R*_0_ and stress 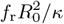 under different Gaussian curvature moduli (a) 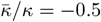 and (b) 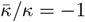.

### C. Determination of Gaussian curvature modulus and spontaneous curvature

In this section, we propose a method to determine the Gaussian curvature modulus 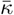 and the spontaneous curvature *c*_0_ simultaneously via a buckling protocol. We assume that the stress-strain relationship of a buckled membrane can be obtained either via molecular dynamics simulation or experiments. In Fig. 6, we show the critical buckling stress *f*_r_(*μ*_r_ = 0) and the stress at half strain *f*_r_(*μ*_r_ = 0.5) as a function of the scaled spontaneous curvature *c*_0_*R*_0_ and the Gaussian bending modulus 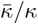. It is found that the contours of the stress are approximately linear, and the slope of the contours are different for *μ*_r_ = 0 and *μ*_r_ = 0.5. Therefore, by measuring the buckled stress at *μ*_r_ = 0 and *μ*_r_ = 0.5, we can determine two lines of different slope, which intersects at one point on the 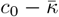 plane, giving the value of the measured 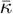 and *c*_0_. The accuracy of the measurement depends on the bandwidth between different contours with a narrow band corresponding to a low accuracy. In order to increase the accuracy, we can use the stress diagram of additional strains. In principle, the Gaussian curvature modulus 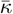 and the spontaneous curvature *c*_0_ can be determined with a very high accuracy when multiple points on the stress-strain relationship are used.

**FIG. 6.**
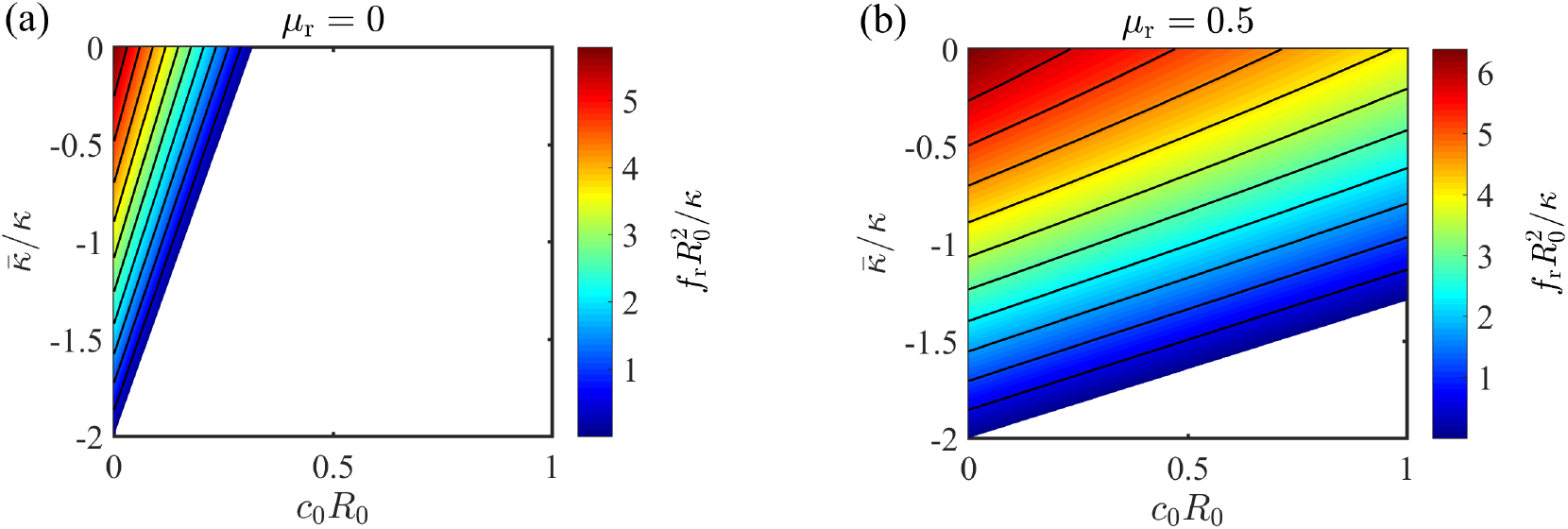
Stress 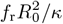 as a function of the scaled spontaneous curvature *c*_0_*R*_0_ and Gaussian curvature modulus 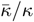 under different strains (a) *μ*_r_ = 0 and (b) *μ*_r_ = 0.5.

## IV. CONCLUSION

In this paper, we aim to study the role of Gaussian curvature modulus 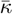 and spontaneous curvature *c*_0_ in membrane buckling under the free-hinge boundary condition by investigating the buckling behaviors of a circular membrane under radially oriented compressive stress. When *c*_0_ and 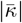 are small, in the absence of the stress, the circular membrane typically adopts a spherical cap shape, which remains unchanged until the stress exceeds a critical value. Then the membrane would buckle in response to the stress and the base radius is reduced to almost zero with increasing stress. A more negative Gaussian curvature 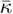 is able to reduce the critical buckling stress to negative values, such that buckling would occur spontaneously in the absence of compressive stresses.

We have proposed a method to determine the Gaussian curvature modulus and the spontaneous curvature simultaneously for molecular dynamics simulations and experiments. The method uses the stress-strain relationship of a circular membrane and compare the measured stress with the calculated stress at different strains. The challenge of this method for molecular dynamics simulations is to impose the free-hinge boundary condition which can not be fulfilled by the typical periodic boundary condition in the simulation box. Implementing this method in molecular dynamics simulations will be our future work.

## ACKNOWLEDGMENTS

We acknowledge financial support from National Natural Science Foundation of China under Grant No. 12174323 and No. 12474199, Fundamental Research Funds for Central Universities of China under Grant No. 20720240144 (RM),and 111 project B16029.

## Appendix A: Derivation of the membrane shape equations

In this section, we provide the mathematical details of how to derive the equations we used to solve the buckled membrane shape. The effective free energy 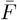 in Eq. (7) is a functional of the membrane shape. The integrand ℒ inside the integral can be written as

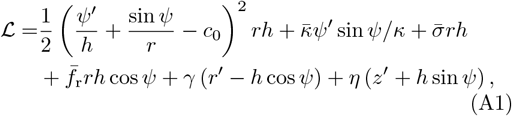

Where 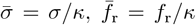, and *γ*(*u*) and *η*(*u*) are Lagrangian multipliers to enforce the geometric relations in Eq. (2). By applying the variational method to the functional 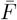, we obtain

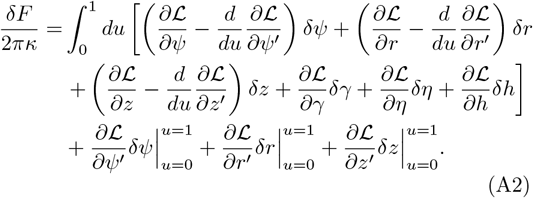

If the bulk terms of Eq. (A2) vanish, we obtain the following shape equations

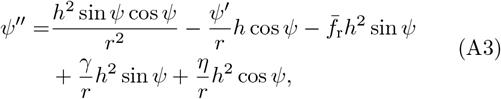

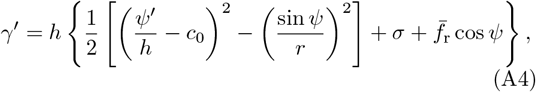

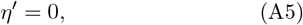

as well as the two geometric relations in Eq. (2). Note that by letting *∂*ℒ*/∂h* = 0, instead of obtaining a shape equation, we derive a conserved quantity

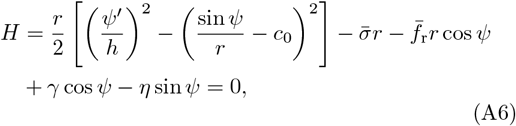

which will be used as a boundary condition.

## Appendix B: Derivation of the boundary conditions

The boundary conditions can be derived by setting the boundary terms in Eq. (A2) to be zero. In particular, at the membrane tip *u* = 0, we fix the angle and the radius by requiring

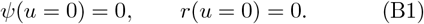

As for the boundary term about *δz*, we let the conjugate term *∂*ℒ*/∂z*^′^ = 0, which means

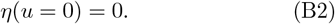

Meanwhile we also require *H*(0) = 0, together with (B1) and (B2), implying that

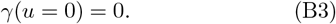

At the membrane base *u* = 1, we impose *∂*ℒ*/∂ψ*^′^ = 0, which gives the free-hinge boundary condition Eq. (8). It can be explicitly written as

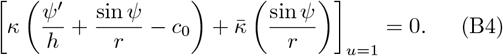

We also impose *δL/δr*^′^ = 0, which is equivalent to

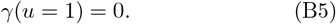

This means that the base radius *r*(*u* = 1) is determined by the equations provided the compressive stress *f*_r_ is given. As for the boundary term about *δz*, we impose *δz*(*u* = 1) = 0, which is to fix the *z*-coordinate

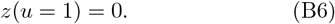

Finally, we impose the incompressibility condition by requiring that the membrane area is a constant

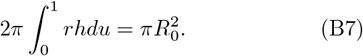

This is achieved by introducing the Lagrangian multiplier *σ* in Eq. (6), and a function *a*(*u*) that satisfies the equation

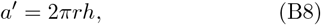

with two boundary conditions

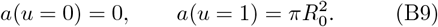

In summary, Eqs. (2), (A3), (A4), (A5) and (B8) make up the full set of shape equations for the buckling problem. They are equivalent to 7 first order ordinary differential equations. In addition, there are two unknown parameters *h* and *σ*, which represent the total arclength and the membrane tension, respectively. Together with the 9 boundary conditions given by Eqs. (B1)-(B6), and (B9), we construct a well-defined boundary value problem.

## Appendix C Numerical method to calculate the stree-strain relationship

In order to obtain the stress-strain relationship and the corresponding buckled membrane shapes, the boundary value problem can be numerically solved with a given stress *f*_r_. We use the boundary value problem solver bvp5c in MATLAB to solve the problem. However, the solver requires a guess of the solution which makes it difficult to obtain the correct solution for an arbitrary value of *f*_r_ if the guessed solution is far from the correct solution. To overcome this issue, we make the stress *f*_r_ an unknown parameter and introduce another boundary condition

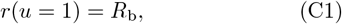

to fix the base radius. We always start from *R*_b_ ≈ *R*_0_ such that the membrane shape is almost flat and the solution is easy to guess. Then we iteratively solve the problem by reducing the base radius *R*_b_ with a small increment Δ*R*_b_ until *R*_b_ is almost zero. The solution for *R*_b_ + Δ*R*_b_ is used for the guessed solution for *R*_b_. In this way, we obtain the stress *f*_r_ for different strain 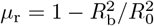, i.e., the stress-strain relationship.

